# Prefrontal High Gamma in ECoG tags periodicity of musical rhythms in perception and imagination

**DOI:** 10.1101/784991

**Authors:** S. A. Herff, C. Herff, A. J. Milne, G. D. Johnson, J. J. Shih, D. J. Krusienski

## Abstract

Rhythmic auditory stimuli are known to elicit matching activity patterns in neural populations. Furthermore, recent research has established the particular importance of high-gamma brain activity in auditory processing by showing its involvement in auditory phrase segmentation and envelope-tracking. Here, we use electrocorticographic (ECoG) recordings from eight human listeners, to see whether periodicities in high-gamma activity track the periodicities in the envelope of musical rhythms during rhythm perception and imagination. Rhythm imagination was elicited by instructing participants to imagine the rhythm to continue during pauses of several repetitions. To identify electrodes whose periodicities in high-gamma activity track the periodicities in the musical rhythms, we compute the correlation between the autocorrelations (ACC) of both the musical rhythms and the neural signals. A condition in which participants listened to white noise was used to establish a baseline. High-gamma autocorrelations in auditory areas in the superior temporal gyrus and in frontal areas on both hemispheres significantly matched the autocorrelation of the musical rhythms. Overall, numerous significant electrodes are observed on the right hemisphere. Of particular interest is a large cluster of electrodes in the right prefrontal cortex that is active during both rhythm perception and imagination. This indicates conscious processing of the rhythms’ structure as opposed to mere auditory phenomena. The ACC approach clearly highlights that high-gamma activity measured from cortical electrodes tracks both attended and imagined rhythms.

---

Neural populations match their activity patterns in response to repetitive, rhythmic auditory stimuli (Nozaradan, 2014; Nozaradan, Peretz, Missal, & Mouraux, 2011; Nozaradan, Peretz, & Mouraux, 2012; Nozaradan, Zerouali, Peretz, & Mouraux, 2015). However, the neural response to rhythmical stimuli is not exclusively driven by exogenous stimulus properties, such as an auditory stimulus, but is also shaped by endogenous top-down mechanisms, such as attention and imagination (Nozaradan et al., 2011). This suggests that neural activity in the context of repetitive auditory stimuli is not only worth investigating as a reactive process, triggered by external stimulation, but may also shed light on complex functions, such as imagination. Here, we aim to further characterise neural activity in auditory perception and imagination, specifically in high-gamma oscillation.

## High Gamma

Recent medical advances have allowed music perception research investigating neural responses to auditory rhythms to venture beyond non-invasive EEG methodologies to invasive measurements such as the use of intracranial electrodes in epilepsy patients. In an intracranial study, Nozaradan, Mouraux, et al. (2017) showed that a 0-30 Hz as well as a 30-100 Hz power band track the envelope of musical rhythms. In the present study, we aim to further explore the involvement of a different power band; High Gamma.

Activity in the high-gamma band is much more localized (Miller et al., 2007) and thought to resemble ensemble spiking (Ray, Crone, Niebur, Franaszczuk, & Hsiao, 2008). Due to the small size of the generator area, frequencies above 70 Hz, become increasingly unreliable to measure, let alone localise, using EEG. In Electrocorticography (ECoG), the electrodes are deployed directly on the cortex rather than on the scalp. This enables accurate characterization of High Gamma (or Broadband Gamma, ~70-170 Hz). This is important, as High Gamma can be linked to auditory attention, auditory perception, and appears to mark auditory segment boundaries (Leuthardt et al., 2011; Pei et al., 2011; Potes, Gunduz, Brunner, & Schalk, 2012; Schalk & Leuthardt, 2011; Sturm, Blankertz, Potes, Schalk, & Curio, 2014)(see Cervenka, Nagle, & Boatman-Reich, 2011 for a review). Indeed, high-gamma activity can even be used to decode speech from the brain (Angrick, Herff, Johnson, et al., 2019; Angrick, Herff, Mugler, et al., 2019; Anumanchipalli, Chartier, & Chang, 2019; Herff et al., 2019; Herff et al., 2015; Pasley et al., 2012). When listening to music, High Gamma averaged across listeners correlates with the sound envelope of a musical piece in a data set with 7 participants (Potes et al., 2012). Using the same data set with an additional three participants, Sturm et al. (2014) found a correlation between High Gamma and the music envelope in 4 out of 10 participants. A recent study also suggests that high-gamma activity is not only involved in music listening, but also music imagination (Ding et al., 2019). In this study, participants were asked to imagination the continuation of familiar musical pieces. High-gamma activity significantly exceeded the baseline that was measured prior to stimulus onset. Using lagged correlations between High Gamma and the music’s envelope, the authors investigated the time course of the activation of different brain regions.

In the present study, we aim to further investigate the potential involvement of High Gamma in music perception. However, rather than exploring familiar musical pieces, we focus on High Gamma’s involvement in musical rhythm perception as well as imagination. Here, we are also less concerned with the time-course of different brain region’s activation, but rather explore areas that capture the underlying periodicities of the rhythmic signal.

## Periodicity tagging

In the present study, we utilise a periodicity tagging approach, which uses autocorrelation representations of the data. This approach focuses on capturing and comparing the periodicities in the musical rhythm with periodicities in high-gamma activity. This approach is inspired by, and related to, the widely used frequency-tagging approach, however, instead of comparing frequency components in the rhythmic envelope with frequency components in neural responses, it compares their periodicities (Henry, Herrmann, & Grahn, 2017; Lenc, Keller, Varlet, & Nozaradan, 2018, 2019; Novembre & Iannetti, 2018; Nozaradan, Keller, Rossion, & Mouraux, 2017; V. G Rajendran, Harper, Garcia-Lazaro, Lesica, & Schnupp, 2017; V. G. Rajendran & Schnupp, 2019). For example, a rhythm might have many interonset intervals (where the onsets are not necessarily consecutive) of 500ms and only a few such interonset intervals of 250ms. Neural responses stimulated by such rhythms might, or might not, exhibit similar temporal periodicities. Because autocorrelation captures the distribution of such periodicities in a signal, measuring the correlation between the autocorrelation of a rhythmic envelope and the autocorrelation of a neural response, allows us to straightforwardly quantify how similar their periodicity distributions are. Autocorrelation is invariant to phase and so is not affected if there is a temporal delay between the two signals. Furthermore, there are a variety of different envelopes that can produce equivalent autocorrelations: we see this as an advantage because it is agnostic to the precise mechanism by which the periodicity is “coded” by the neural envelope. Indeed, there are various ways in which high-gamma activity could code the stimulus – not only through envelope matching – so a many-to-one matching is necessary when looking for areas of interest that track periodicity of stimuli. Here, we argue that if a high-gamma brain activity pattern represents or tracks the underlying periodicity of an acoustic signal, then it is most likely related to the stimulus. In summary, we specifically ask here whether high-gamma oscillations during listening, as well as imagination of repetitive auditory rhythms captures the rhythms periodicities using a periodicity tagging (autocorrelation) approach.

## Method

### Participants

Electrocorticographic data were recorded from 8 patients (3 female, 5 male, 22 to 42 years old) with pharmacoresistant epilepsy undergoing localization of epileptogenic zones and eloquent cortex before surgical resection. When questioned, no patients reported hearing deficits or any form of musical training. In all cases, a tumour was not the source for the seizures and no lesions were indicated by any electrode used for analysis. Patients participating in this study gave written informed consent and the study protocol was approved by the institutional review boards of Old Dominion University and Mayo Clinic, Florida. Patients were implanted with subdural electrode grids or strips, based purely on their clinical need. Electrode locations were verified by co-registering pre-operative MRI and post-operative CT. For combined visualization, electrode locations were projected to common Talairach space. Electrode locations and activations were rendered using NeuralAct (Kubanek & Schalk, 2015). We recorded electrocorticographic activity during rhythm perception and imagination of a total of 437 (151 left hemisphere, 286 right) subdural electrodes (see Figure 1).

### Stimuli

The majority of research investigating neural activity to auditory rhythms stimuli use either complex speech or simple clicks, white noise, pure tones, or sine tones (Nozaradan, Mouraux, et al., 2017). To increase ecologically validity for musical stimuli, we use kick-snare drum patterns. Here, we analyse data of participants listening to two different musical rhythms. Each rhythm consists of 8 pulses and 4 sounded events. The rhythms are being presented at either 120 bpm or 140 bpm. Table 1 presents a summary of all rhythms. Rhythm 2 is a syncopated rhythm, that is listeners will perceive a downbeat on the 5^th^ element, despite there being no sounded event. We included a syncopated rhythm, as syncopation is typically considered to increase rhythmic complexity (Fitch & Rosenfeld, 2007); this allows us to explore periodicity tagging in a more complex rhythm. Furthermore, a control was implemented by a condition that presented white noise instead of a rhythm.

**Table 1.** Overview of the musical rhythms

| Rhythm | Sounded Events | Sequence |  |  |  |  |  |  |  |
| --- | --- | --- | --- | --- | --- | --- | --- | --- | --- |
| Unsyncopa | 4 | K | x | S | x | K | x | S | x |
| Syncopatec | 4 | K | x | S | K | x | S | x | x |

*Note.* ‘K’ represents a sounded Kick, ‘S’ represent a sounded Snare, and ‘x’ represent a non-sounded element.

### Procedure

Each participant passively listened to each rhythm in condition-blocks of 8 repetitions in 120 bpm and 10 repetitions in 140 bpm. Within each rhythm block, the rhythm dropped out (i.e., became silent) for two repetitions in the 120 bpm condition and two repetitions during the 140 bpm condition and participants were instructed to imagine the rhythm to continue (imagined condition). Each block appeared twice throughout the experiment. The order of rhythm blocks was randomised. For the listening and imagination blocks, participants were instructed not to tap along with the rhythm or to move, and adherence to these instructions was confirmed for each participant through investigator observation. For each rhythm block, additional trials were performed that required participant to tap the events of the rhythms using their dominant hand. These intermingled tapping trials as well as additional rhythm blocks using different rhythms were not included in the present analysis. ECoG signals were simultaneously recorded throughout the experiment.

### ECoG data collection

Data from the electrode grids or strips Ad-Tech Medical Instrument Corporation, Wisconsin, 1 cm spacing) were band-pass filtered between 0.5 and 500 Hz and recorded using g. USB amplifiers (g.tec, Austria) at a sampling rat of 1200 Hz. Data recording and stimulus presentation were facilitated by BCI2000 (Schalk, McFarland, Hinterberger, Birbaumer, & Wolpaw, 2004). Electrode grids for all eight participants can be seen in Figure 1.

**Figure 1.**
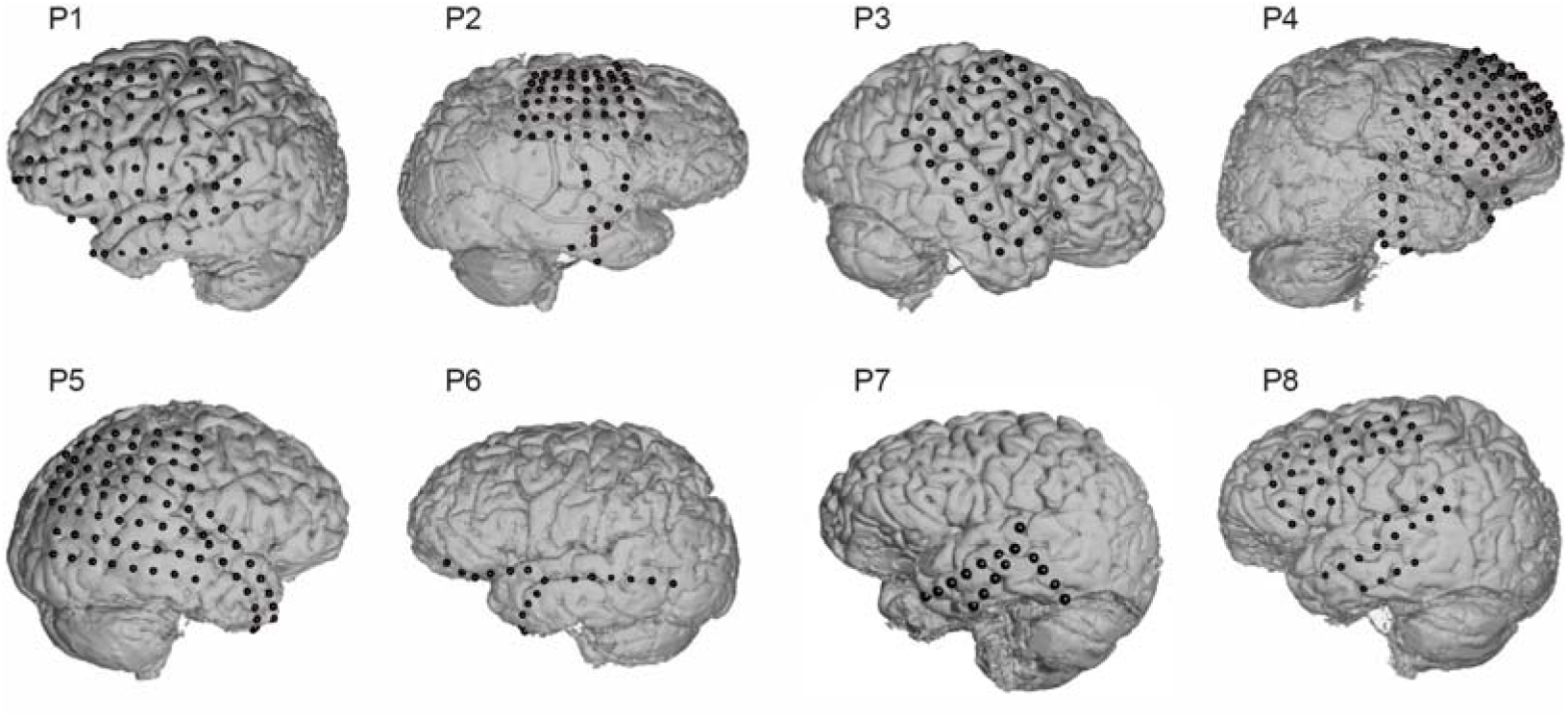
Electrode grid locations for all eight participants.

### Data Analysis

Separately for each participant, electrode, tempo (120 bpm vs. 140 bpm), audio condition (listening vs imagine), and rhythm (Unsyncopated: K x S x K x S x vs. Syncopated: K x S K x S xx) we extracted the absolute Hilbert envelope of High-gamma activity. We used elliptic IIR low- and high-pass filters to bandpass filter the ECoG signals between 70 and 170 Hz and applied an elliptic IIR notch filter to attenuate the first harmonic of the 60 Hz line noise. The Hilbert transform was then used to extract the envelope. We calculated the circular autocorrelation over all repeated presentations of the rhythm up to the Nyquist frequency. This was done by taking the real component of an inverse DFT of a pointwise multiplication of a DFT of the High Gamma timeseries and its complex conjugate and then dividing each element by the maximum element of the vector. The same transformation was conducted on the envelope of the musical rhythm’s waveform. High Gamma and musical rhythm autocorrelations were correlated with one another. The resulting autocorrelation correlations (*ACC*) between high-gamma brain activity and musical rhythms were used to statistically assess whether High Gamma tracks musical rhythms. This process is schematically represented in Figure 2. Visually, ACC can be described as the correlation between the top and bottom right panels in Figure 2. As a control, we extracted high-gamma activity, envelope and autocorrelation also for a condition where participants were listening to white noise instead of the actual musical rhythms and calculated ACC.

**Figure 2.**
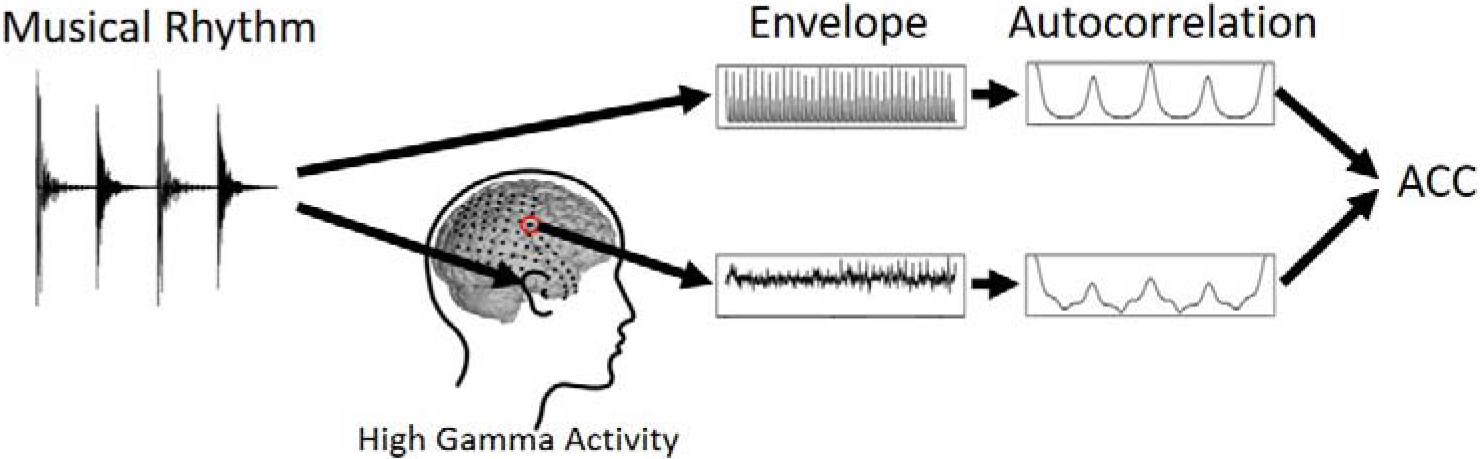
Schematic representation of the data analysis. The left most panel shows the original waveform of a musical rhythm. The rhythm in this example is a ‘K x S x K x S x’, with ‘K‘ being the kick, ‘S’ the snare, and ‘x’ a pause. First, we extracted the envelope of the continuously looped presentation of the rhythm, as shown in the middle top panel. The top right panel then shows the autocorrelation of the rhythm’s envelope. Note that the shown autocorrelation vector corresponds to the length of the original rhythm to emphasise the relationship between waveform and autocorrelation. For the actual analysis we used the whole autocorrelation vector over all repeated presentations of the rhythm up to the Nqyuist frequency. Simultaneously, we measured high-gamma activity from cortex electrodes whilst participants are listening (or imagining) the rhythm. Similar to the musical rhythm, we extracted the envelope of the high-gamma activity and calculated the autocorrelation. In a last step, we correlated the autocorrelations of high-gamma envelope and musical rhythm envelope to obtain our dependent variable: autocorrelation correlation (ACC).

We deployed a Bayesian mixed effect model predicting the correlation between the autocorrelations of High-Gamma and musical rhythms (*ACC*, scaled to *Mean* = 0, *SD* = 1) based on *Rhythm* (Unsyncopated. vs. Syncopated), and *AudioCondition* (listen vs. imagine), and *Signal* (white noise vs. rhythm). The model was provided with a random effect for *Participant*, *Electrode*, *Tempo* (120bpm vs. 140bpm), and *Presentation* (first vs second time a condition was shown), resulting in the maximal random effect structure as justified by the experimental design (Barr, Levy, Scheepers, & Tily, 2013). The models were implemented in the R-environment (R-Core-Team, 2013) using the brms-package (Bürkner, 2017, 2018). The *Signal* coefficient in combination with its interaction terms allows us to inspect the evidence in favour of whether autocorrelation in high-gamma oscillation meaningfully tracks musical rhythms, whilst controlling for brain activity that a participant – at a given electrode location – would show when listening to a length-matched white noise segment instead of the actual musical rhythm. In other words, our model is provided with the information of how high the ACCs value between High Gamma and musical rhythms can be expected to be, simply because participants are listening to any auditory noise (here we use white noise as a control) individually for every electrode of each participants. The model predicts the difference to this baseline, when participants are actually listening or imagining the rhythms. The model was provided with a weakly informative prior student-*t*(3, 0, 1), and ran on 4 chains with 1000 warm-ups and 10000 iterations each.

## Results

In a first step, we explore whether High-Gamma tracking periodicities of the musical rhythms can be observed on a broad spatial scale. For this, we deployed Bayesian mixed effects models that compare ACCs obtained when participants listened or imagined the rhythms to ACCs obtained from the baseline. The baseline is ACC between a musical rhythm and High Gamma of a given participant and electrode when listening to white noise. Table 2 shows coefficient estimates (*β)*, 95%-CIs, as well as *Evidence Ratios* for the hypothesis that there is elevated brain-wide High Gamma tracks the musical rhythm. For convenience, we denote with * conditions that show ‘significant’ tracking of periodicities at an *α* = .05 level (*Evidence Ratios* > 19) (Milne & Herff, accepted 2020). The results of table 2 are derived by performing the hypothesis tests shown in table 3 on the fitted model shown in table 4.

**Table 2.** Summary of evidence observed in each condition whether broad spatial High Gamma tracks the periodicities of musical rhythms more than baseline.

| Rhythm | Audio Condition | $\beta$ | 95%-CI$_{\beta}$ | Evidence Ratio |
| --- | --- | --- | --- | --- |
| Unsyncopated | Listen | .1136 | .0827 to .1445 | > 9999* |
| Unsyncopated | Imagine | -.0435 | -.0749 to -.0125 | .0114 |
| Syncopated | Listen | .0741 | .0432 to .1052 | > 9999* |
| Syncopated | Imagine | .1368 | .1049 to .1678 | > 9999* |

*Note.* We obtain strong evidence for broad spatial High Gamma tracking of the envelope of musical rhythms in the syncopated rhythms during listening and imagining, as well as in the unsyncopated rhythm during imagining. However, we do not observe evidence for whole-brain tracking of the unsyncopated rhythm during. \* denote effects that can be considered ‘significant’ at a $\alpha = .05$ level.

**Table 3.** Hypotheses performed on the model shown in table 4.

| Rhythm | Audio Condition | Hypothesis test |
| --- | --- | --- |
| Unsyncopated | Listen | Rhythm > 0 |
| Unsyncopated | Imagine | Rhythm + Rhythm:Imagined > 0 |
| Syncopated | Listen | Rhythm + Rhythm:Syncopated > 0 |
| Syncopated | Imagine | Rhythm + Rhythm:Syncopated + Rhythm:Imagined + Rhythm:Syncopated:Imagined > 0 |

*Note.* The reference was placed at the white noise condition, unsyncopated, listening condition. *Rhythm* indicates that the actual rhythm rather than white noise was heard. *Unsyncopated* and *Syncopated* refer to the two different rhythms used. *Imagined* indicates that the rhythms was not played, and instead participants were asked to imagine them.

**Table 4.** Model summary.

| *Predictors* | **Scaled AC Correlation** |  |
| --- | --- | --- |
|  | *Estimates* | *CI (95%)* |
| Intercept | -0.67 | -0.73 to -0.61 |
| Rhythm | 0.11 | 0.08 to 0.15 |
| Syncopated | 0.44 | 0.40 to 0.48 |
| Imagined | 0.82 | 0.78 to 0.85 |
| Rhythm.Syncopated | -0.04 | -0.09 to 0.01 |
| Rhythm.Imagined | -0.16 | -0.21 to -0.11 |
| Syncopated.Imagined | -0.00 | -0.05 to 0.05 |
| Rhythm.Syncopated.Imagined | 0.22 | 0.15 to 0.29 |
| N<sub>Electrode</sub> | 518 |  |
| Observations | 16576 |  |
| Marginal R<sup>2</sup> / Conditional R<sup>2</sup> | 0.216 / 0.629 |  |
| •<sup>2</sup> | 0.41 |  |

On a broad spatial scale, we observe strong evidence (all *Evidence Ratios* > 9999) in favour of high-gamma ACs tracking the ACs of the musical rhythms in the syncopated rhythm during listening and imagination, and in the unsyncopated rhythm during listening, but not imagination. When comparing the two rhythms, the unsyncopated and the syncopated rhythms show comparable ACCs in the listening condition (*β* = .04, *EE* = .03, 95%-CI_β_ = -.004 to .83, *Evidence Ratio* = 13.46). In the imagination condition, however, we obtain strong evidence for higher ACCs in the syncopated rhythm compared to the unsyncopated condition (*β* = .18, *EE* = .03, 95%-CI_β_ = .14 to .22, *Evidence Ratio* = >9999*). Although we do not observe tracking on a broad spatial scale in the unsyncopated imagination condition, this does not imply that there are no electrodes for which the high-gamma oscillation tracks the musical rhythms, as can be seen in the electrode-wise results.

Figure 3 shows counts of the electrodes that significantly track the musical rhythms’ periodicities, as well as their normalised ACC. We calculated significance thresholds for each participant and rhythm individually. For this, we used the distribution of correlations between the autocorrelation of a musical rhythm and the autocorrelation of high-gamma activity whilst listening to length-matched white noise segments. Correlations that exceed 99% of this distribution are deemed significant. Normalised ACC values were obtained by subtracting the ACC when listening to length-matched white noise instead of listening or imagining the musical rhythms. Each electrode in a given participant was normalised by the white noise ACC of the same electrode in that participant. As can be seen in Figure 3, each condition contains electrode in which high-gamma ACs track the ACs of the respective musical rhythms.

**Figure 3.**
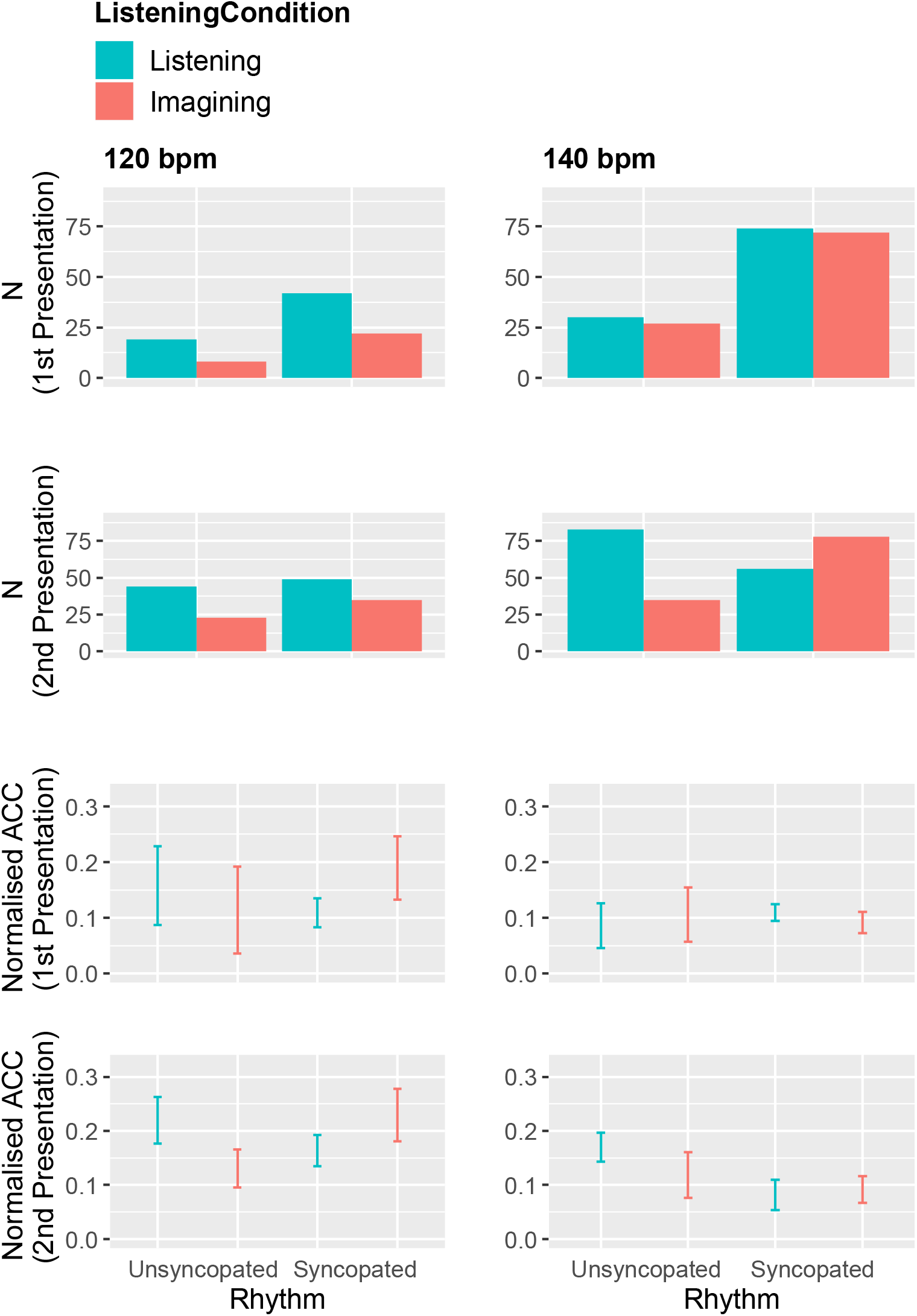
Number (first and second row) and magnitudes (third and fourth row) of electrodes that significantly track musical rhythms in their high-gamma activity, pooled across participants and electrodes. Significance was defined by exceeding participant-wise 99% of the ACC between musical rhythms and High Gamma during the white-noise control condition. All conditions contain electrodes that significantly track the musical rhythms. Normalised ACC values were obtained by subtracting the significance thresholds from the observed ACCs. Error bars represent 95%-CIs.

Figure 3 also suggests an increase in the number of significant electrodes between first and second presentation of each condition (i.g., higher bars in the second row compared to the first). A Bayesian mixed model supports this. The model predicts *Normalised ACC* based on *Presentation Number* (first vs. second), whilst controlling for *Participant* and *Electrode*. The model reveals an increase in Normalised ACC in all conditions (all *Evidence Ratios* > 65*). This can be seen in Table 5.

**Table 5.**
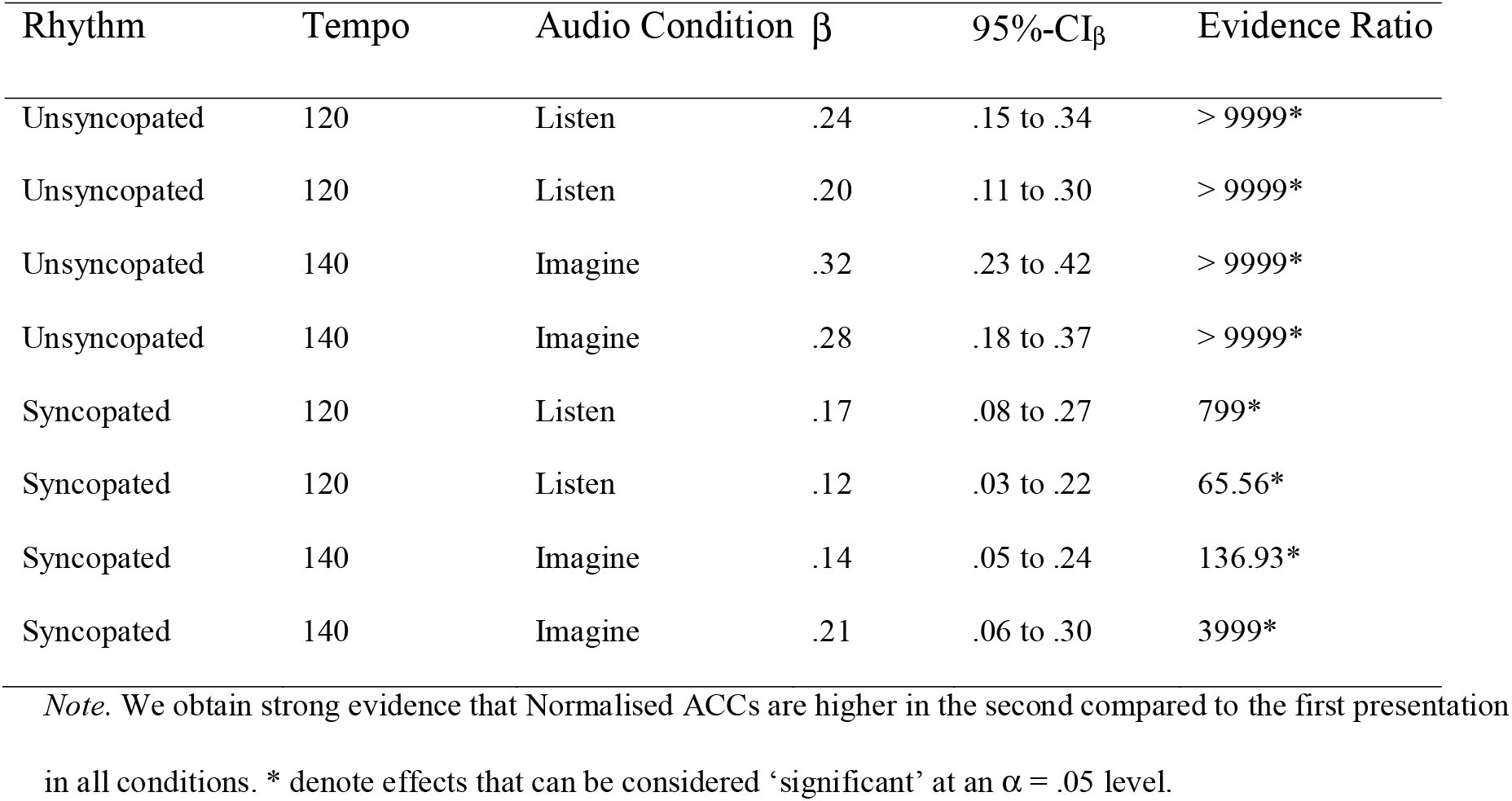
Summary of evidence observed that Normalised ACCs are higher during the second presentation compared to the first presentation of each condition.

| Rhythm | Tempo | Audio Condition | $\beta$ | 95%-CI$_{\beta}$ | Evidence Ratio |
| --- | --- | --- | --- | --- | --- |
| Unsyncopated | 120 | Listen | .24 | .15 to .34 | > 9999* |
| Unsyncopated | 120 | Listen | .20 | .11 to .30 | > 9999* |
| Unsyncopated | 140 | Imagine | .32 | .23 to .42 | > 9999* |
| Unsyncopated | 140 | Imagine | .28 | .18 to .37 | > 9999* |
| Syncopated | 120 | Listen | .17 | .08 to .27 | 799* |
| Syncopated | 120 | Listen | .12 | .03 to .22 | 65.56* |
| Syncopated | 140 | Imagine | .14 | .05 to .24 | 136.93* |
| Syncopated | 140 | Imagine | .21 | .06 to .30 | 3999* |

*Note.* We obtain strong evidence that Normalised ACCs are higher in the second compared to the first presentation in all conditions. \* denote effects that can be considered ‘significant’ at an $\alpha = .05$ level.

To investigate the potential overlap between significant electrodes in listening and imagination we used a Bayesian Mixed effects models predicting *SignificanceDuringImagination* (Binary factor with 1 = significant, 0 = not significant), based on *SignificanceDuringListening* (and vice versa), whilst controlling for *Rhythm, Tempo, Participant, Presentation*, and *Electrode*. We observe very strong evidence that *SignificanceDuringListening* predicts *SignificanceDuringImagination* (*β* = 1.83, *EE* = .25, 95%- CI_β_ = 1.43 to 2.24, *Evidence Ratio* = >9999*) and vice versa (*β* = 2.51, *EE* = .26, *95%-CI_β_* = 2.01 to 2.94 *Evidence Ratio* = >9999*). This suggests high predictive information between the electrodes that are significant in listening and those that are significant during imagination. Further insight is provided in the topography section of the results.

To visualise the tracking, Figure 4 shows examples for each condition. The red line shows the AC of a given musical rhythm. The blue line shows the AC of an example electrode.

**Figure 4.**
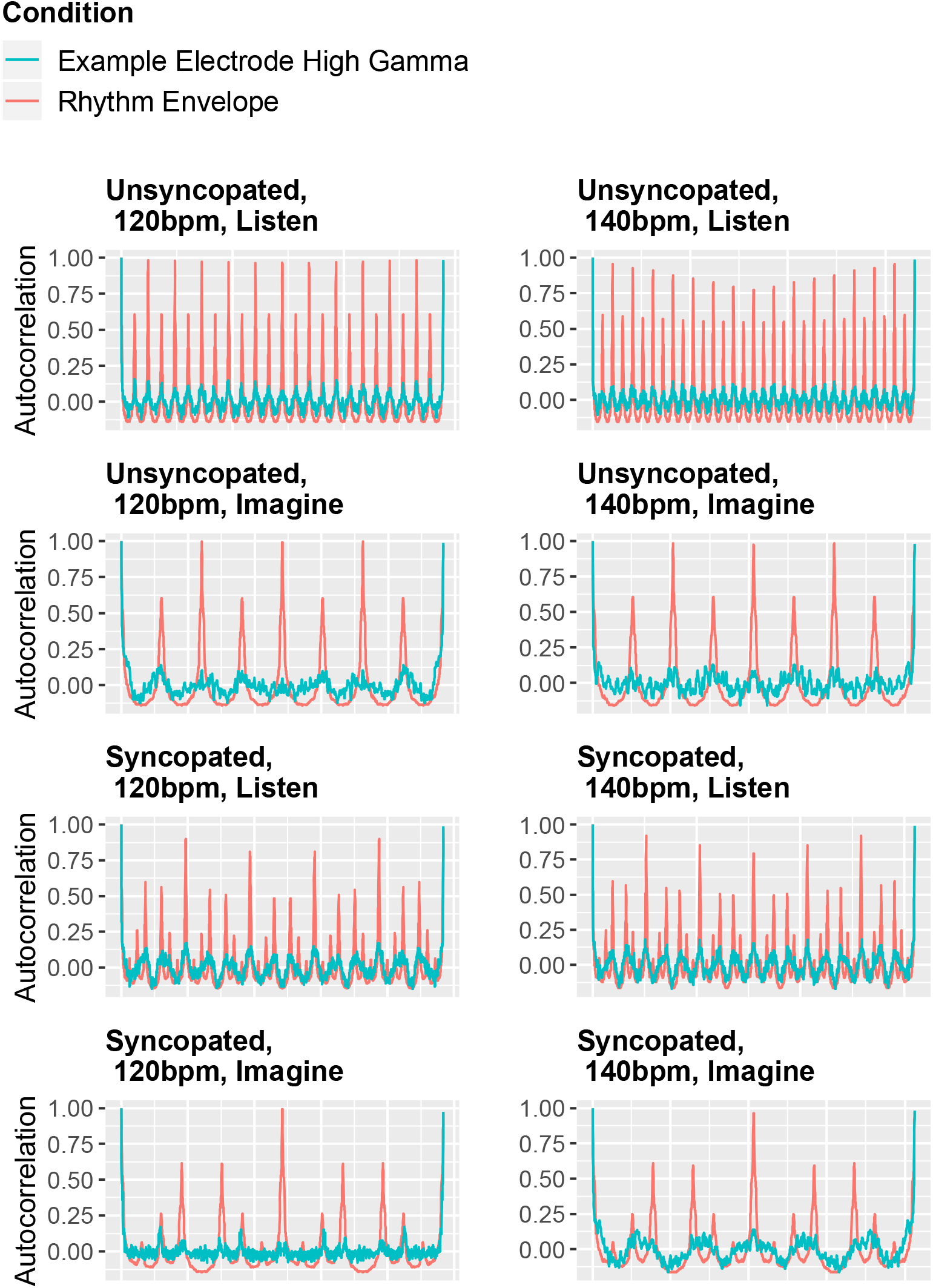
ACs of the musical rhythm conditions and example electrodes. There are electrodes in which High Gamma ACs (blue) significantly track the musical rhythms ACs (red) in all conditions.

This study is predominantly concerned with High Gamma, however, we performed the same analysis on the Beta band (12-30 Hz) to see whether high-gamma oscillation carries information that is not contained in other frequency bands. We chose Beta because it was suggested by the reviewers and prior work suggests an involvement of Beta in neural processing of musical rhythms (Chang, Bosnyak, & Trainor, 2016). We observed strong evidence that there are more electrodes that significantly track the musical rhythm’s AC using High Gamma compared to Beta (*β* = 2.17, *EE* = .72, *95%-CI_β_* = 0.98 to 3.4, *Evidence Ratio* = 2799*). Furthermore, the increase in Normalised ACC between first and second presentation that is observed in all conditions in High Gamma is not observed in Beta in any condition (all Evidence Ratios < 5.78), with the exception of the unsycopated rhythm at 140bpm in the imagined condition (*Evidence Ratio* = 799*). However, it is worth mentioning that we also found some electrodes that tracked the musical rhythms’ ACs in the Beta ACs.

### Topography

To localise the effect, we plotted all electrodes on a joint brain map. Figure 5 shows heat maps of mean normalized ACC for listening (top) and imagining (bottom) across all rhythms and tempi.

**Figure 5.**
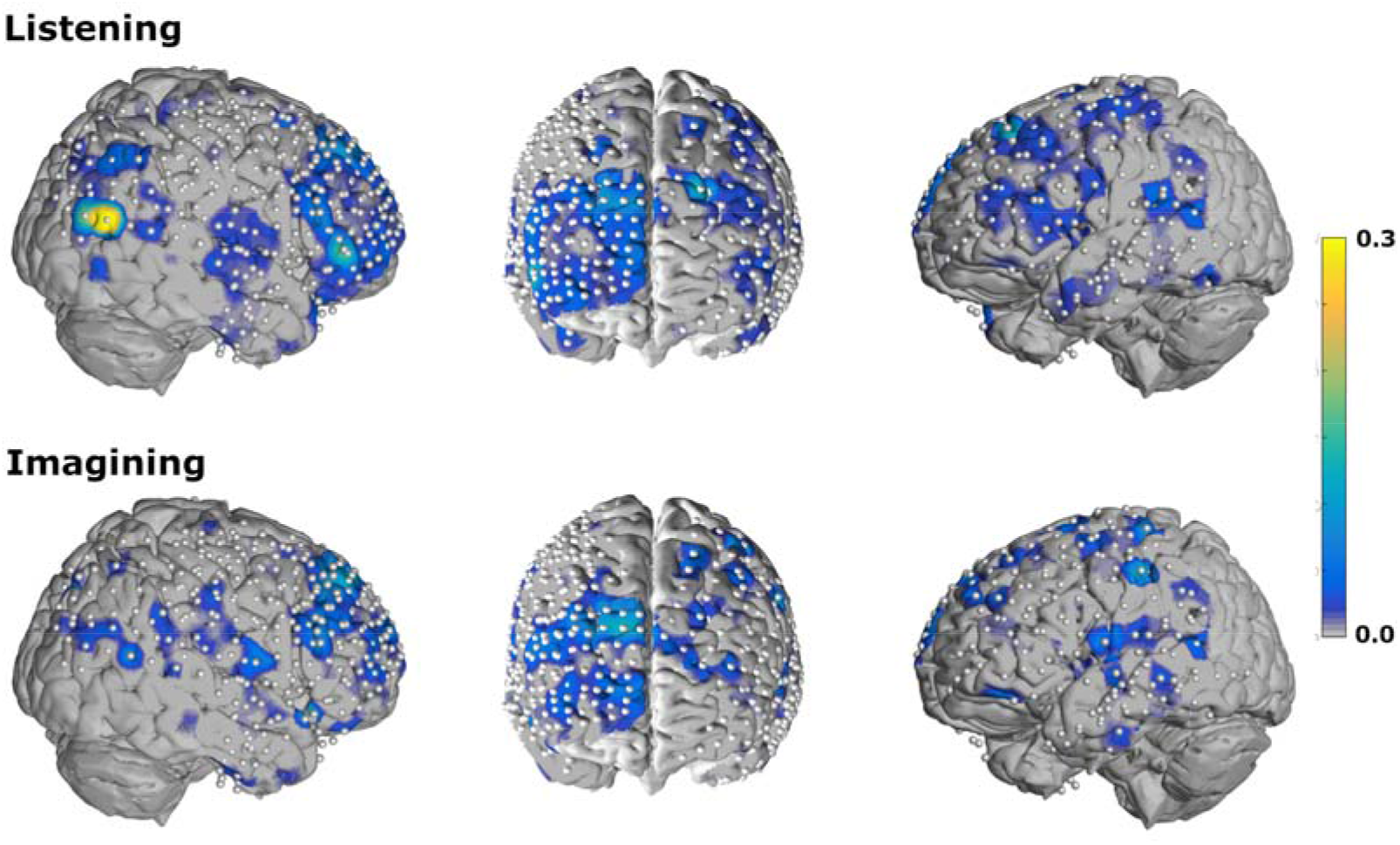
A joint brain map for all participants across all conditions. Heat maps visualize mean normalized ACC across all rhythms and tempo. Significant ACCs can be observed particularly in the frontal areas of the right hemisphere. These ACCs are also significant during the imagine condition.

## Discussion

The present study investigated the involvement of High Gamma in listening as well as imagining musical rhythms using brain activity of nine participants measured through invasive ECoG. Bayesian mixed effects models provided compelling support that high-gamma oscillation tracks the envelope of musical rhythms. Specifically, we deployed an analytical approach that emphasises the periodicity in musical rhythms by investigating correlations between the autocorrelations of musical rhythms’ and the autocorrelation of high-gamma brain activity. In all listening conditions the models support the conclusion that high-gamma activity captures the periodicity in musical rhythms. We observe the same in all but one condition, when participants are imagining the rhythms, rather than listening to them. Taken together, it appears that during imagination, neural populations display similar high-gamma activity that tracks the envelope of the imagined stimulus, usually observed when acoustic stimuli are actually present. This may be preliminary support for the notion that – on a neural level – imagination involves activity of the reactive neural response associated with the presence of the stimulus.

The present finding supports previous ECoG studies that highlight the importance of high-gamma activity in auditory processing (Herff et al., 2015; Leuthardt et al., 2011; Pasley et al., 2012; Pei et al., 2011; Potes et al., 2012; Schalk & Leuthardt, 2011; Sturm et al., 2014) (see Cervenka et al., 2011). Specifically, our results replicate the findings that High Gamma tracks music envelopes (Sturm et al., 2014). Such replications are important, because ECoG studies operate with very small sample sizes. Furthermore, we extend the finding to imagination, and a periodicity tagging approach. The direct approach of directly correlating High Gamma with stimulus envelope deployed by (Sturm et al., 2014) relies on relatively long segments, clean data, and a phase-locking. Furthermore, correlating High Gamma with the stimulus envelope can only identify neural population that engage in envelope matching. Yet, there are various ways in which high-gamma activity could theoretically code the stimulus. The present approach is able to identify neural populations that engage in envelope matching as well as those that match any form of distinct activity pattern to the periodicities of the stimuli. As such, we put the present approach forward as a useful tool to identify brain regions of interest. The identified regions could then be further analysed to characterise the nature of the activity pattern that tracks the periodicities of the stimuli. It is important to note that the present approach correlates the two autocorrelations with one another. It is possible that other metrics of similarity such as cosine similarity, Weissman score, or shared mass could work equally well or even better. Future work could investigate the benefits of more sophisticated similarity measures.

The unsyncopated and the syncopated rhythms show comparable ACCs in the listening condition. This is worth noting as the syncopated rhythm could be considered the more complicated rhythm (Fitch & Rosenfeld, 2007). The stronger tagging of the unsyncopated rhythm compared to the syncopated rhythm in the imagination condition is surprising. A possible explanation could be, that the syncopated rhythm may be more interesting and engaging for participants, having a ‘groove’ that makes it easier to entrain. A different explanation considering the order in which the conditions were presented is provided in the limitation section.

High-gamma activity showed a greater number of significant electrodes compared to beta activity. High Gamma also shows a strong increase in Normalised ACCs between first and second presentation in all conditions. This increase was only seen in one condition for Beta (Unsyncopated, 140bpm, imagined). The strong evidence for an increase in Normalised ACCs between first and second presentation in High Gamma, but not Beta, may suggest that some form of higher order auditory processing is involvement in periodicity tagging in High Gamma that improves with increased exposure. A possible candidate could be a prediction-based mechanism that shows clearer activation patterns when familiar with a rhythm. While High Gamma showed more significant electrodes, there were some electrodes that also showed significant tagging of the musical rhythms’ periodicities in the Beta band. This is interesting because Beta can be reliable captured in EEG, whereas High Gamma cannot. A future study could investigate whether periodicity tagging can be shown using EEG and the autocorrelations in the Beta band.

### Topography

Electrodes with significant ACCs can be found in auditory areas in the superior temporal gyrus and in frontal areas on both hemispheres. Numerous significant electrodes are observed on the right hemisphere which is in accordance with previous findings (Thaut, Trimarchi, & Parsons, 2014). Of particular interest is the large cluster of electrodes in the right prefrontal cortex that are active during both rhythm perception and imagined perception, which indicate conscious processing of the rhythm structure as opposed to mere auditory phenomena. This finding mirrors research that also observed frontal High Gamma when imagining familiar music (Ding et al., 2019). The previous study also found elevated high-gamma activation in the temporal lobe during imagination. Here, we did not observe that High Gamma in the temporal lobe represents the periodicities of the musical rhythms during imagination like the prefrontal cortex does. However, this could simply be due to a difference in methodology. The previous study (Ding et al., 2019) focused on areas that show elevated high-gamma activity and/or areas where gamma activity tracks the music’s envelope. The present study uses musical rhythms rather than familiar music, and focuses on areas that track the rhythms’ periodicities, regardless of overall activity. However, any area that closely tracks the audio envelope in the present dataset would have been identified by our periodicity-tagging approach, so further research is required to shed line of the role of temporal lobe during imagination.

### Limitations

An important limitation in the present design is that what we and others (Ding et al., 2019) liberally term ‘imagination’ is in fact an ‘imaginary continuation’ of the rhythms. In theory, such a continuation could be functionally distinct from un-prompted imagination. In fact, it is possible that if the imagination condition would have lasted longer, then potentially the high-gamma representation of the rhythms’ periodicity may have diverged. This is an empirical question for a future study. Furthermore, the one condition that did not show brain-wide significant tracking of the rhythms periodicity in High Gamma urges caution interpretation of the present results. This is, because the condition was the simpler rhythm. If anything, we would have expected this condition to show the strongest effect. A possible explanation lies in the fact that this condition was always tested first. Potentially, participants were not yet familiar with the imagination task to evoke a reliable effect. Some support for this explanation can be gained from the increase in Normalised ACC as well as number of significant electrodes between first and second presentation of the conditions. Furthermore, as common in invasive brain studies, we were operating with small participant numbers, and despite our best efforts of making the most of the data at hand, by deploying a Bayesian framework, we simply may not have the statistical power to compensate for all sources of random variability.

## Conclusion

Deploying an analytical approach that emphasises the periodicity in musical rhythms, we found that high-gamma brain activity in auditory areas tracks periodicity when listening to musical rhythms. Furthermore, we found that high-gamma activity in the prefontal cortex tracks periodicity of musical rhythms both during listening and imagination.

## Acknowledgements

We would like to thank all participants for their participation and immense contribution. This research was supported by two grants of the National Science Foundation (NSF 1451028, NSF 1902395).

## Author Contributions

SAH wrote the manuscript and performed the statistical analysis. CH, CDJ, and SAH preprocessed the ECoG data. AJM developed the periodicity tagging approach. SAH implemented the periodicity tagging approach and adjusted it to the present study. DJK and GDJ designed the experiment and recorded the data. JJS contributed to the patient recruitment, data collection, and clinical aspect of the study. CH, AJM, GDJ, JJS, and DJK provided input on the manuscript.

## Significance Statement

The possibility to capture high-frequency brain activity, such as High Gamma, with high spatial and temporal resolution makes invasive brain recordings extremely valuable. We present new data from an invasive ECoG study with a comparably large sample size. Deploying a new periodicity-tagging technique that extends the common frequency-tagging, we found that High Gamma in auditory areas tracks periodicity. Furthermore, we use the periodic nature of musical-stimuli as a neural footprint and found that high-gamma activity in the prefontal cortex tracks periodicities of musical rhythms both during listening and imagination. The neural mechanisms involved in imagination in particular are ill understood. The present study provides evidence that the pre-frontal cortex tracks periodicities in auditory stimuli during perception and imagination, and highlights the usefulness of musical stimuli for studying neural processes.

